# Production of ORF8 protein from SARS-CoV-2 using an inducible virus-mediated expression system in suspension-cultured tobacco BY-2 cells

**DOI:** 10.1101/2020.10.07.325910

**Authors:** Tomohiro Imamura, Noriyoshi Isozumi, Yasuki Higashimura, Shinya Ohki, Masashi Mori

## Abstract

COVID-19, caused by severe acute respiratory syndrome coronavirus 2 (SARS-CoV-2), which spread worldwide in 2020, is an urgent problem to be overcome. The ORF8 of SARS-CoV-2 has been suggested to be associated with the symptoms of COVID-19, according to reports of clinical studies. However, little is known about the function of ORF8. As one of the ways to advance the functional analysis of ORF8, mass production of ORF8 with the correct three-dimensional structure is necessary. In this study, we attempted to produce ORF8 protein by chemically inducible protein production system using tobacco BY-2 cells. An ORF8-producing line was generated by the *Agrobacterium* method. As a result, the production of ORF8 of 8.8 ± 1.4 mg/L of culture medium was confirmed. SDS-PAGE and nuclear magnetic resonance (NMR) analysis confirmed that the ORF8 produced by this system is a dimeric form with a single structure, unlike that produced in *Escherichia coli*. Furthermore, it was suggested that the ORF8 produced by this system was glycosylated. Through this study, we succeeded in producing ORF8 folded into a single structure in a chemically inducible protein production system using tobacco BY-2 cells. It is expected that the functional analysis of ORF8 will be advanced using the ORF8 produced by this system and that it will greatly contribute to the development of antibodies and therapeutic agents targeting ORF8.

---

Coronavirus disease 2019 (COVID-19), caused by severe acute respiratory syndrome coronavirus 2 (SARS-CoV-2) and characterized by severe pneumonia, spread worldwide in 2020. To overcome the threat of SARS-CoV-2, the development of therapeutic agents and vaccines based on functional elucidation is urgently needed. The SARS-CoV-2 genome was sequenced recently (Acc. No.: NC_045512.2, Fig. 1a). Interestingly, open reading frame 8 (ORF8), an accessory protein of SARS-CoV-2, has lower homology than that of the closely related SARS virus, unlike other SARS-CoV-2 proteins. Instead, ORF8 of SARS-CoV-2 shows high homology with ORF8 of bat coronavirus (Lu, et al. 2020). SARS-CoV-2 mutants lacking ORF8 were reported to be less likely to cause severe disease (Young, et al. 2020). ORF8 is reported to be highly immunogenic, and anti-ORF8 antibodies are formed in the early stage of infection (Hachim, et al. 2020). Furthermore, it has been reported that a significant T-cell response to ORF8 is observed in recovered patients (Grifoni, et al. 2020). Thus, ORF8 is predicted to be closely associated with the symptoms caused by SARS-CoV-2. An X-ray crystallization study has revealed the three-dimensional structure of ORF8 (Flower, et al. 2020) and that ORF8 affects the immune system (Zhang, et al. 2020). Further studies on ORF8 function are warranted.

**Fig. 1.**
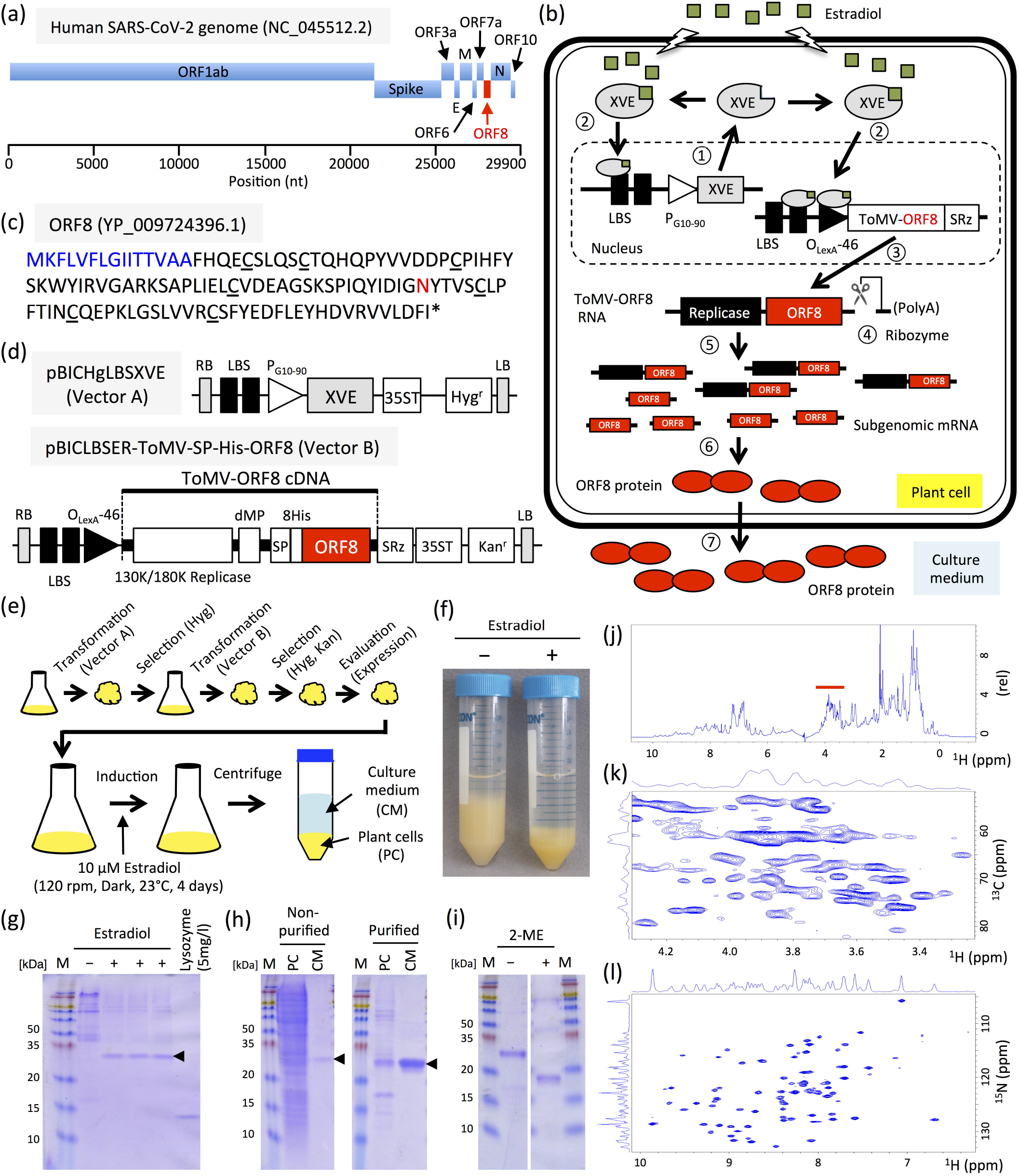
Production and characterization of ORF8. (a) Schematic representation of SARS-CoV-2 genome. E, M, and N indicate envelope protein, membrane glycoprotein, and nucleocapsid phosphoprotein, respectively. (b) Schematic overview of the chemically inducible ToMV-mediated expression system. Step 1: Constitutive expression of XVE, a transcriptional activator. Step 2: XVE binds to estradiol and is activated. Step 3: Activated XVE binds to O_LexA-46_ promoter or LBS to promote transgene transcription. Step 4: Non-viral sequences that interfere with the replication of the viral vector are removed by ribozymes. Step 5: Viral vector replication. Step 6: Translation of ORF8 from subgenomic RNA. Step 7: Translocation of ORF8. (c) Amino acid sequence of ORF8. Blue and red letters indicate signal peptide and *N*-glycosylation site, respectively. Underbars indicate cysteine. (d) Schematic representation of ToMV-mediated chemically induced expression plasmids. XVE, transcriptional activator that responds to estrogen; SP, signal peptide of *Arabidopsis* chitinase; 8 His, 8× His-tag; LBS, LexA binding site; P_G10-90_, synthetic constitutive promoter; P_LexA-46_, fusion promoter controlled by XVE; dMP, partial movement protein; SRz, ribozyme sequence from tobacco ringspot virus satellite RNA; 35ST, 35S terminator; RB, right border; and LB, left border. Hyg^r^ and Kan^r^ indicate expression cassette of hygromycin and kanamycin resistance genes, respectively. (e) Generation of ORF8-producing cell line and induction of ORF8 production. (f) Four days after 17β-estradiol induction of suspension-cultured tobacco BY-2 cells; + and − indicate with or without 17β-estradiol, respectively. (g) Detection of ORF8 from 7.5 μL of cultured medium by Coomassie Brilliant Blue staining. + and − indicate with or without 17β-estradiol, respectively. Arrowhead indicates ORF8. Lysozyme (5 mg/L) indicates loading control. (h) Non-purified and purified ORF8. PC and CM indicate crude extracts of plant cells and culture medium, respectively. (i) Molecular property of ORF8. + and − indicate with or without 2-mercaptoethanol (2ME), respectively. M, protein size marker. Numbers indicate molecular weights (kDa). (j) ^1^H NMR spectra of unlabeled ORF8. Red line indicates the glycan signal appearing region. (k) ^1^H-^13^C HSQC of unlabeled ORF8 protein. An expanded view of the glycan signal appearing region is shown. (l) ^1^H-^15^N HSQC of ^15^N-labeled ORF8.

To analyze ORF8 function, a large amount of homogeneous protein folded into a single structure needs to be produced, which can also be used for developing antibodies and therapeutic agents to treat COVID-19. A plant-based production system can be used to mass produce eukaryotic proteins that are physiologically active. We have developed a highly efficient chemically inducible protein production system using tobacco BY-2 cells, utilizing the extremely high protein production capacity of tobamovirus (ToMV) (Dohi, et al. 2006). In this study, we used an improved expression system that provided positive feedback on transcription factor (XVE) expression (Fig. 1b). Protein production using plant-cultured cells is a large-scale process that is uniform, aseptic, and GMP-compliant. Our system can mass produce eukaryotic proteins with multiple disulfide bonds with sufficient quality for structural and functional studies. We have successfully produced proteins that were difficult to produce uniformly in *Escherichia coli.* including plant-bioactive (Costa, et al. 2014; Ohki, et al. 2011) and animal (Ohki, et al. 2008) proteins and have determined their structures using solution nuclear magnetic resonance (NMR). Mature ORF8 contains seven cysteines, six of which form three intramolecular disulfide bond pairs and one forms an intermolecular disulfide bond for dimerization (Flower, et al. 2020) (Fig. 1c). In addition, one *N*-glycosylation site has been predicted for ORF8 (Fig 1c). Our production system is expected to be suitable for uniform ORF8 protein production with these structural features.

To produce ORF8, we designed the amino acid sequence for the 8 His-tagged ORF8 fusion protein with an extracellular signal peptide added to the N-terminus. Next, artificial *ORF8* was synthesized by optimizing codons in tobacco and introducing restriction enzyme sites for cloning at both ends (IDT, Coralville, USA; Acc. No.: LC586256). The artificially synthesized *ORF8* was introduced into a chemically inducible tobamovirus vector (pBICLBSER-ToMV-SP-His-ORF8, Fig. 1d). Next, pBICLBSER-ToMV-SP-His-ORF8 and pBICHgLBSXVE (Fig. 1d) expressing the artificial transcription factor XVE, which activates transcription by binding with 17β-estradiol (Dohi, et al. 2006), were introduced into tobacco BY-2 cells by the *Agrobacterium* method, and the ORF8-producing line was selected (Fig. 1e). The ORF8-producing cells were suspension-cultured in normal MS medium and MS medium labeled with an ^15^N nitrogen source for NMR analysis. ORF8 production was induced by adding 10 μM 17β-estradiol (Fig. 1e). After 4 days, the induced tobacco BY-2 cells had significantly suppressed proliferation due to viral vector replication compared to non-induced cells (Fig. 1f). ORF8 production was confirmed in the induced cells. ORF8 was the major protein found in the culture medium (Fig. 1g). This production system could produce 8.8 ± 1.4 mg of ORF8 per liter culture medium. Following purification using a nickel column, ORF8 in the culture medium could be obtained in the form of a single protein (Fig. 1h). In *E. coli*, ORF8 did not form the correctly folded three-dimensional structure and required refolding; moreover, a mixture of monomers and dimers was produced (Flower, et al. 2020). In contrast, the molecular weight of ORF8 produced by tobacco BY-2 cells was changed by approximately 50% in the presence of 2-mercaptoethanol (2ME) compared with that in the absence of 2ME. This result indicates that ORF8 produced in tobacco BY-2 cells are dimers due to the formation of disulfide bonds (Fig. 1i). Furthermore, ^1^H and ^1^H-^13^C HSQC NMR analysis suggested that ORF8 was glycosylated (Fig. 1j, k), indicating that ORF8 is glycosylated in infected individuals. Moreover, distribution, line-shape, and intensity of the peaks in ^1^H and ^1^H-^15^N HSQC strongly indicated that ORF8 produced using the BY-2 system forms a stable β-sheet-rich three-dimensional structure (Fig. 1j, l). The number of peaks in ^1^H-^15^N HSQC was less than expected, probably because the larger molecular size due to glycosylation and dimerization caused line broadening.

In this study, we successfully produced ORF8 that is folded into a single structure in a ToMV-mediated chemically induced protein production system using tobacco BY-2 cells. Furthermore, the purification process could be simplified by releasing ORF8 into the culture medium that was low in plant-derived contaminants. Our finding shows that it is possible to produce ORF8 folded into a single structure in large quantities with high efficiency. Because ORF8 has an intact three-dimensional structure, it can be used for functional analysis and the production of antibodies and therapeutic agents targeting ORF8.

## Acknowledgments

This work was partly supported by the Nanotechnology Platform of MEXT, Japan. The authors thank Akiko Mizuno, Hiroko Hayashi, and Akio Miyazato for technical assistance.

## Author contributions

MM conceived this study. MM and TI designed the experiments. All authors performed the experiments. TI, SO, and MM wrote the manuscript. All authors have read and approved the final manuscript.

